# Genetic disruption of leucine rich repeat transmembrane protein 4 like 1 induces a pro-social behavioural phenotype in zebrafish

**DOI:** 10.1101/2025.05.12.653402

**Authors:** Courtney Hillman, Giulia Petracco, Barbara D Fontana, María Florencia Scaia, Elisa Dalla-Vecchia, Jon H Wetton, William HJ Norton, Matthew Parker, Florian Reichmann

**Author notes:** Corresponding authors: Dr. Florian Reichmann, Division of Pharmacology, Otto Loewi Research Center, Medical University of Graz, Neue Stiftingtalstraße 6, A-8010 Graz, Austria, Dr Matthew Parker, Surrey Sleep Research Centre, University of Surrey, Clinical Research Building, Egerton Road, Guildford, GU2 7XP, UK.

## Abstract

**Background:** Social behaviour encompasses the wide range of interactions that occur between members of the same species. In humans, disruptions in social behaviour are characteristic of many neuropsychiatric disorders, where both genetic risk factors and synaptic dysfunctions can contribute to the phenotype. Among the genes implicated in synaptic regulation, the synaptic adhesion protein leucine-rich repeat transmembrane protein 4 (LRRTM4) has been identified as a key player in maintaining synaptic function and neuronal circuit integrity. Despite its established role in the nervous system, the potential involvement of LRRTM4 in modulating social behaviour and its contribution to social deficits has yet to be explored.

**Methods:** In the current study, we used zebrafish to study how genetic deletion of *lrrtm4l1*, a zebrafish orthologue of *LRRTM4*, affects sociality. For this, the social behaviour of homozygous *lrrtm4l1* knockout (KO) zebrafish was analysed in multiple behavioural assays and the brain transcriptome of mutant animals was investigated by RNAseq.

**Results:** KO zebrafish displayed a pro-social phenotype in multiple behavioural assays. Groups of *lrrtm4l1* KO zebrafish formed more cohesive shoals and KO individuals spent more time in the vicinity of conspecifics during a social interaction test. They were also less aggressive and in contrast to wild-type zebrafish did not differentiate in their interactions with known and unknown groups of fish. Neurotranscriptomic analysis revealed 560 differentially expressed genes including changes in glutamatergic neurotransmitter signalling, tryptophan- kynurenine metabolism and synaptic plasticity.

**Conclusion:** These findings suggest that *lrrtm4l1* is an important regulator of social behaviour in zebrafish. In a translational perspective, LRRTM4 is a promising potential therapeutic target that warrants further investigation in the framework of neuropsychiatric conditions characterized by major social impairments.

## Introduction

Social behaviour refers to the broad spectrum of potential interactions between individuals of a species. Sociality is common among animals, likely reflecting survival benefits, such as forming swarms or shoals to deter potential predators (1). Social interactions are also the basis for reproductive success ensuring the survival of the species, which is deeply evolutionary rooted (2). Among humans, the nature and quality of social relationships are fundamental for wellbeing and mental health (3). However, throughout the progression of neuropsychiatric disorders, social behaviour is frequently disrupted, presenting a significant challenge for the affected individual and their immediate social environment (4). Numerous factors have been recognised as key contributors to the emergence of social disturbances, such as genetic makeup, as well as brain circuit, neuronal and synaptic dysfunctions (5-7).

Synaptic adhesion proteins are important regulators of proper synaptic functioning and neuronal circuit integrity. Leucine rich repeat transmembrane proteins (LRRTMs) are a subclass of this protein family with four known members in the human genome (8). They are all expressed at neuronal postsynaptic membranes across the central nervous system exhibiting different expression patterns and functions. All members of this protein family contain so-called leucine-rich repeats, which enable them to form trans-synaptic contacts with various presynaptic binding partners (9). The binding of LRRTM4 to presynaptic glypicans regulates hippocampal synapse development, presynaptic differentiation and proper functioning of retinal GABAergic synapses in mammals (10-12). LRRTM4 has also been implicated in glutamatergic neurotransmission, through the regulation of alpha-amino-3- hydroxy-5-methyl-4-isoxazole propionic acid (AMPA) receptor trafficking at postsynaptic sites (13). *Lrrtm4* knockout mice show decreased dendritic spine density of hippocampal dentate gyrus neurons and *Lrrtm4* knockdown in cultured cortical neurons leads to reduced dendritic and PSD-95-positive spine density (13,14)

At the synaptic level, LRRTMs have been studied extensively, but evidence about their role during behaviour is sparse. In the context of social behaviour, only LRRTM1 has been studied so far. Specifically, *Lrrtm1*^*−/−*^ mice have been found to display a claustrophobia-like phenotype, but they also spent more time interacting with an intruder mouse in the resident-intruder task (15). In contrast, a recent study reported no alterations in the social behaviour of *Lrrtm1*^*−/−*^ mice in the three-chamber social interaction test, but social impairments of *Lrrtm1/SynCAM1* double knockout mice (16). Recent findings from human genetic studies suggest that *LRRTM4* is associated with the development of neuropsychiatric disorders. These disorders include autism spectrum disorder and Tourette syndrome, both of which are often characterised by severe social deficits (17,18). In addition, genetic aberrations at the *LRRTM4* locus have been linked to children’s aggressive behaviour (19) indicating an important role of this protein across multiple social domains.

In the current study, we used zebrafish to study how genetic deletion of *lrrtm4l1*, a zebrafish orthologue of *LRRTM4*, affects social behaviour. Zebrafish display a large spectrum of social behaviours that can be reliably assessed in established behavioural assays (20-22). In addition, zebrafish have been widely used to study the genetic basis of disorders characterised by social deficits (23,24). We and others have previously shown that *lrrtm4l1* is highly expressed in the brain of zebrafish with a similar expression pattern compared to *LRRTM4* expression in humans (25,26). Given that *LRRTM4* polymorphisms have been linked to autism and aggressive behaviour as described above, we performed behavioural and brain transcriptomic analysis of *lrrtm4l1* mutant zebrafish to characterise the role of this gene during social behaviour and to identify biological pathways underlying the behavioural phenotype.

## Results

### Lrrtm4l1 deletion induces a pro-social phenotype

To investigate how *LRRTM4* regulates social behaviour, we generated a novel zebrafish mutant line of the zebrafish *LRRTM4* orthologue, *lrrtm4l1*, using CRISPR/Cas9. The mutation of the novel line is a 1bp insertion located early in exon 2 of the gene, leading to a frame-shift and an early stop codon (Fig.1a). This results in a strongly truncated, likely non-functional protein, given that the whole cytosolic and transmembrane domain of the protein as well as large parts of the extracellular protein domain are not translated (Suppl. Fig.1). To characterise the social phenotype of this mutant line, we first analysed shoaling behaviour. In this assay, 5 fish per genotype are placed together in a large tank to analyse shoal cohesiveness. Strikingly, we found that *lrrtm4l1*^*−/−*^ (KO) shoals had a lower average minimum and maximum distance between the fish of a shoal, a lower average distance between all shoal members and a smaller average shoal area compared to *lrrtm4l1*^*+/+*^ (WT) shoals indicating a pro-social phenotype (Fig.1b). This view is also supported by the behaviour of the KO zebrafish in the visually-mediated social preference test (Norton et al. 2019), where mutants spent more time in the vicinity of conspecifics than corresponding WT fish (Fig.1c). To analyse whether this pro-social behaviour might be related to reduced aggression of the KO fish, we performed the mirror-induced aggression (MIA) assay (27) and found that KO zebrafish were indeed less aggressive (Fig.1d).

**Fig. 1.**
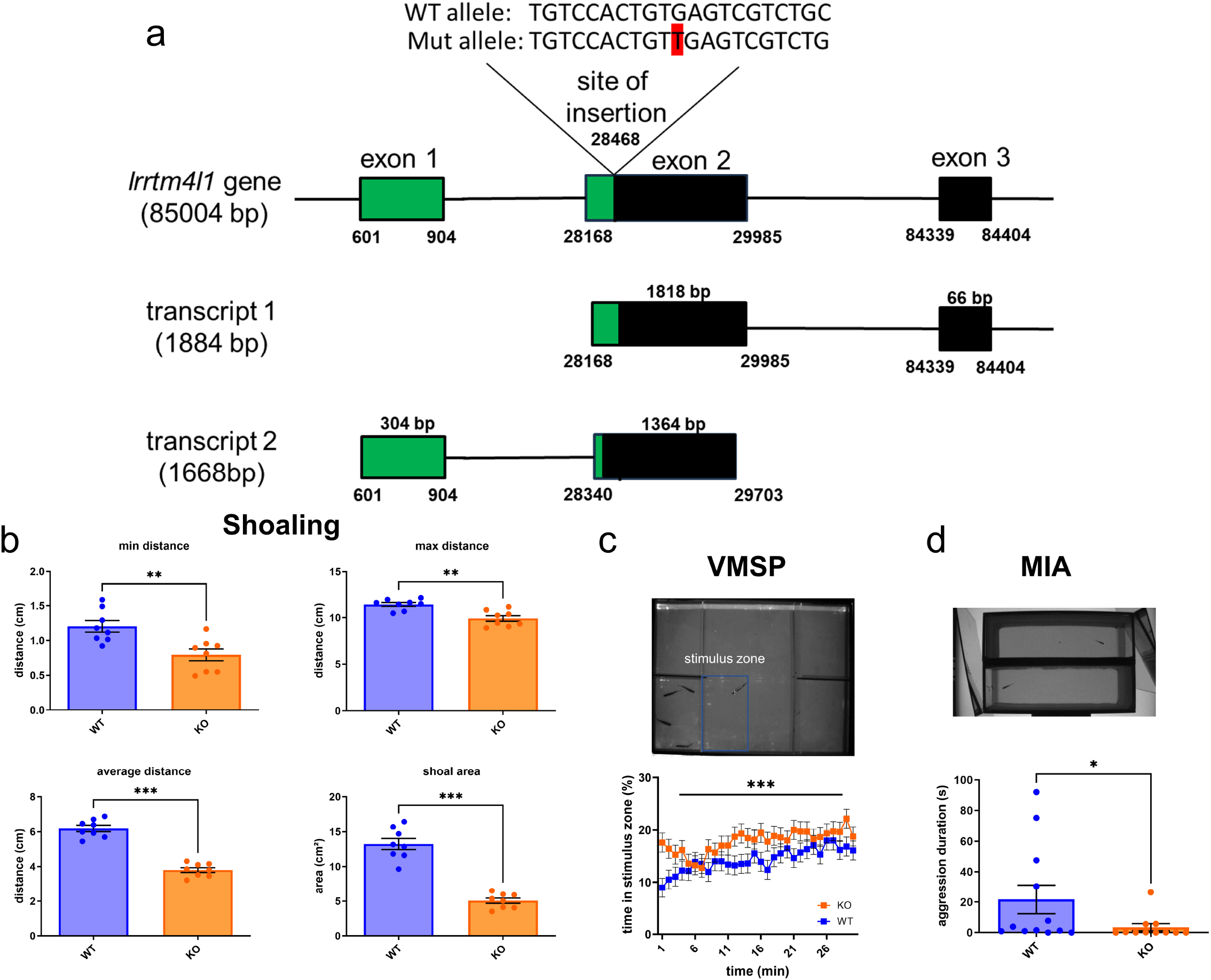
Social phenotype of lrrtm4l1^−/−^ zebrafish. (a) Schematic representation of the lrrtm4l1 gene and its transcripts. The site and type (1bp insertion) of the mutation within the gene is shown at the top. The 1bp insertion is highlighted in red. Green colour indicates the parts of the gene and its transcripts unaffected by the mutation, while the black parts are not translated. (b) Shoaling assay. Lrrtm4l1^−/−^ (KO) zebrafish swim closer together and form a more cohesive shoal than their wild-type (WT) counterparts. Student’s t-test. n = 8 shoals/group. (c) Visually-mediated social preference (VMSP) test. KO zebrafish spend more time in the stimulus zone than respective WT fish. Linear mixed model, between subjects (WT vs KO) effect: P < 0.001 followed by unpaired Student’s t-test. n=12/group. (d) Mirror-induced aggression assay. KO zebrafish are less aggressive against their mirror image. Mann-Whitney U test. n=11-12/group. ^***^P < 0.001; ^**^P < 0.01; ^*^P < 0.05 KO vs WT. Data are presented as mean ± SEM

### *lrrtm4l1*^*−/−*^ zebrafish show dysregulated responses to social context

Given the altered social behaviour of the *lrrtm4l1*^*−/−*^ zebrafish, we next investigated whether social context has an influence on their pro-social phenotype. For this, we analysed the behaviour of KO and WT fish exposed to a known stimulus shoal (from the same home tank) or an unknown stimulus shoal in the visually-mediated social preference (VMSP) test (22). As can be seen in Fig. 2a, WT zebrafish spent more time in the vicinity of an unknown shoal compared to a known shoal, which might be an expression of social curiosity. Mutant zebrafish, however, did not interact more with an unknown shoal, although their overall social interaction time was higher than that of WT zebrafish (Fig. 2b). This might indicate that KO zebrafish are incapable of differentiating between known and unknown zebrafish shoals or that they choose to strongly interact with stimulus shoals independent of prior experience with them. To test the hypothesis that KO zebrafish do not differentiate between known and unknown shoals, we next examined whether their interactions with stimulus shoals involved aggressive components. Notably, WT zebrafish showed reduced aggression in response to an unknown shoal, an effect absent in mutants (Fig. 2c,d), supporting the hypothesis that they fail to differentiate between known and unknown shoals. Finally, we tested whether social isolation of the test fish affects their social interaction behaviour with both known and unknown shoals. Similar to non-isolated WT fish, isolated WT zebrafish spent more time in the vicinity of an unknown shoal than a known shoal (Fig. 2e). Interestingly, however, under these conditions KO zebrafish also show increased sociability towards the unknown shoal (Fig. 2f). This indicates that after strong social challenges such as isolation, *lrrtm4l1*^*−/−*^ zebrafish are capable of social recognition and that this intervention increases their social curiosity towards the unknown shoal.

**Fig. 2.**
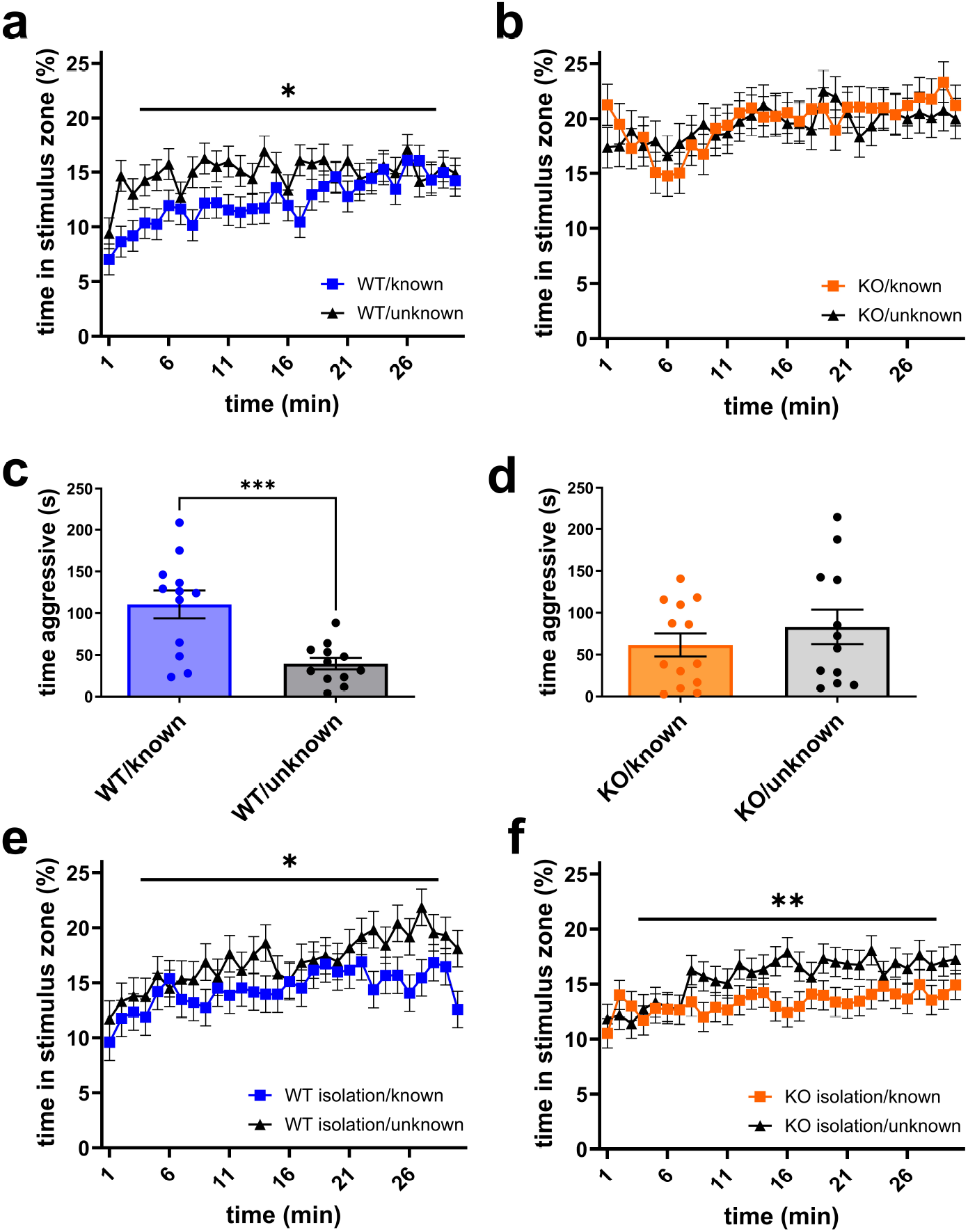
Effects of stimulus shoal characteristics and social isolation on social interaction behaviour. (a) Time spent near conspecifics of wild-type (WT) zebrafish exposed to a group of known or unknown fish in the visually-mediated social preference test (VMSP) analysed with a linear mixed model and an unpaired Student’s t-test. n = 12/group. (b) Time spent near conspecifics of lrrtm4l1^−/−^ (KO) zebrafish exposed to a group of known or unknown fish in the VMSP analysed using a linear mixed model followed by an unpaired Student’s t- test. n = 12/group. (c) Aggressive behaviour of WT zebrafish exposed to a group of known or unknown fish in the VMSP. Student’s t test. n = 12/group. (d) Aggressive behaviour of KO zebrafish exposed to a group of known or unknown fish in the VMSP. Student’s t test. n = 12/group. (e) Time spent near conspecifics of socially isolated WT zebrafish exposed to a group of known or unknown fish in the VMSP. Linear mixed model followed by Student’s t- test. n = 12/group. (f) Time spent near conspecifics of socially isolated KO zebrafish exposed to a group of known or unknown fish in the VMSP. Linear mixed model followed by Student’s t-test. n = 12/group. ^***^P < 0.001; ^**^P < 0.01; ^*^P < 0.05 KO vs WT. Data are presented as mean ± SEM.

### *lrrtm4l1* deletion alters the expression of neurotransmitter-related and synaptic plasticity genes in the zebrafish brain

To study molecular mechanisms underlying the behavioural alterations, we analysed differences in the brain transcriptome between *lrrtm4l1*^*−/−*^ and *lrrtm4l1*^*+/+*^ zebrafish. Principal component analysis (PCA) of the 200 most variable genes revealed two sample clusters clearly separating WT from KO samples along PC1 of the plot (Fig.3a) indicating major transcriptional differences. Differential expression analysis revealed 560 differentially expressed genes (DEGs; padj ≤ 0.05 and LFC ≥ |0.58|) between the genotypes, out of which 322 were upregulated and 238 were downregulated in KO samples (Fig.3b; Suppl. Table 1). The 10 most significant DEGs are involved in diverse biological processes (Fig.3c). Among them, is the gene *cytoplasmic FMRP-interacting protein 2* (*cyfip2*), which is downregulated in KO zebrafish (Fig.3d). This is of interest, because in mice excitatory neuron-specific constitutional knockout of *Cyfip2* has been linked to social dominance behaviour (28). Another key finding is the strong downregulation of *tryptophan 2*,*3-dioxygenase* a (*tdo2a*), a key enzyme in tryptophan metabolism (Fig.3c,e), that catalyses the rate limiting step in the kynurenine pathway. The observed downregulation of *tdo2a* indicates a decrease in the production of kynurenine and thus a shift of tryptophan metabolism towards alternative pathways such as the serotonin pathway (29). Interestingly, we also found that *kynurenine 3- monooxygenase* (*kmo*), which converts kynurenine to 3-hydroxy kynurenine, a cytotoxic compound, is upregulated in mutants (Fig. 3f) suggesting an accumulation of neurotoxic kynurenine metabolites. Given that LRRTM4 has been linked to activity-dependent synaptic insertion of AMPA receptors (13), it is of interest to note that *glutamate receptor, ionotropic, AMPA 2a* (*gria2a*), encoding an AMPA receptor subunit, is downregulated after *lrrtm4l1* KO (Fig.3g). In addition, we detected an upregulation of *glutamate receptor, metabotropic 2a* (*grm2*) in mutants, which encodes a metabotropic glutamate receptor (Fig.3h). Interestingly, we also found differences in potassium channel expression (Fig.3i-k) and in *vesicle amine transport 1* (*vat1*) expression, a gene that encodes a synaptic vesicle protein (Fig.3l) suggesting further alterations in neurotransmitter signalling in the mutants. We also found changes in synaptic plasticity-related genes such as *synaptophysin a* (*sypa*), *glypican 2* (*gpc2*), a family member of the glypicans that have been found as binding partner of LRRTM4 (9) and *semaphorin 3F* (*sema3f*), a critical mediator of axon guidance (Fig. 3m-o; (30)).

**Fig. 3.**
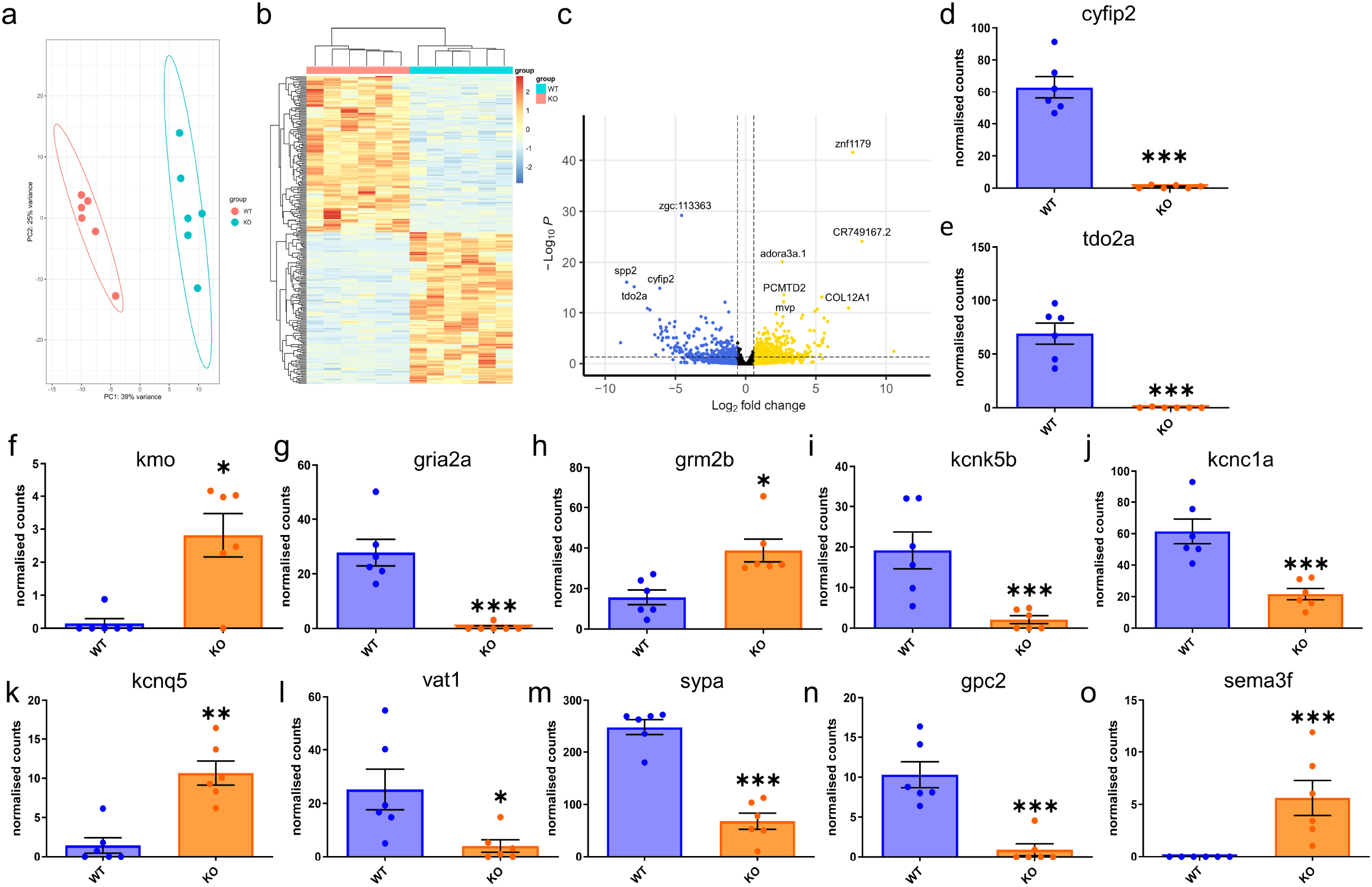
Neurotranscriptomic effects of lrrtm4l1 deletion. a) Principal component analysis plot of the 200 most variable genes after differential expression analysis between lrrtm4l1^−/−^ (KO) and lrrtm4l1^+/+^ (WT) zebrafish. (b) Heatmap of differentially expressed genes (DEGs; padj < 0.05 and LFC > |0.58|) between KO and WT zebrafish. Hierarchical clustering of samples and genes reveals large differences between KO and WT, but similar transcriptional patterns within the two lines. (c) Volcano plot displaying the most significant DEGs between KO and WT zebrafish. Golden dots indicate genes upregulated in KO more than log fold change 0.58, blue dots represent genes downregulated in KO more than LFC –0.58 and black dots represent genes not passing these thresholds as indicated by vertical dotted lines. (d-o) Selected DEGs between WT and KO zebrafish. (d) cytoplasmic FMRP-interacting protein 2 (cyfip2), (e) tryptophan 2,3-dioxygenase a (tdo2a), (f) kynurenine 3- monooxygenase (kmo), (g) glutamate receptor, ionotropic, AMPA 2a (gria2a), (h) glutamate receptor, metabotropic 2a (grm2), (i) potassium channel, subfamily K, member 5b (kcnk5b), (j) potassium voltage-gated channel, Shaw-related subfamily, member 1a (kcnc1a), (k) potassium voltage-gated channel, KQT-like subfamily, member 5a (kcnq5a), (l) vesicle amine transport 1 (vat1), (m) synaptophysin a (sypa), (n) glypican 2 (gpc2), (o) semaphorin 3F (sema3f). n = 6/group. Wald test with Benjamini-Hochberg correction. ^***^P < 0.001; ^**^P < 0.01; ^*^P < 0.05 KO vs WT. Data in d-o are DESeq2-normalised counts presented as mean ± SEM.

To analyse which biological pathways are enriched in our dataset, we used the database for annotation, visualization and integrated discovery (DAVID) (31). As shown in Fig.4a, functional annotation clustering revealed 7 significantly enriched clusters with an enrichment score ≥ 1.3. The most enriched cluster (cluster 1) contains pathway terms related to the lysosome and DEGs encoding lysosomal enzymes, transporters and receptors (Suppl. table 2) suggesting that *lrrtm4l1* KO disturbs lysosomal degradation and/or autophagy. Other clusters found to be enriched contain more general processes such as oxidoreductase activity (cluster 2), ATP/nucleotide binding (cluster 3) or nucleosomal DNA binding (cluster 6). These findings were also confirmed by over-representation analysis using g:profiler (32), which revealed an enrichment of nucleosome binding, nucleosomal DNA binding and lysosomal processes (Fig.4b).

**Fig. 4.**
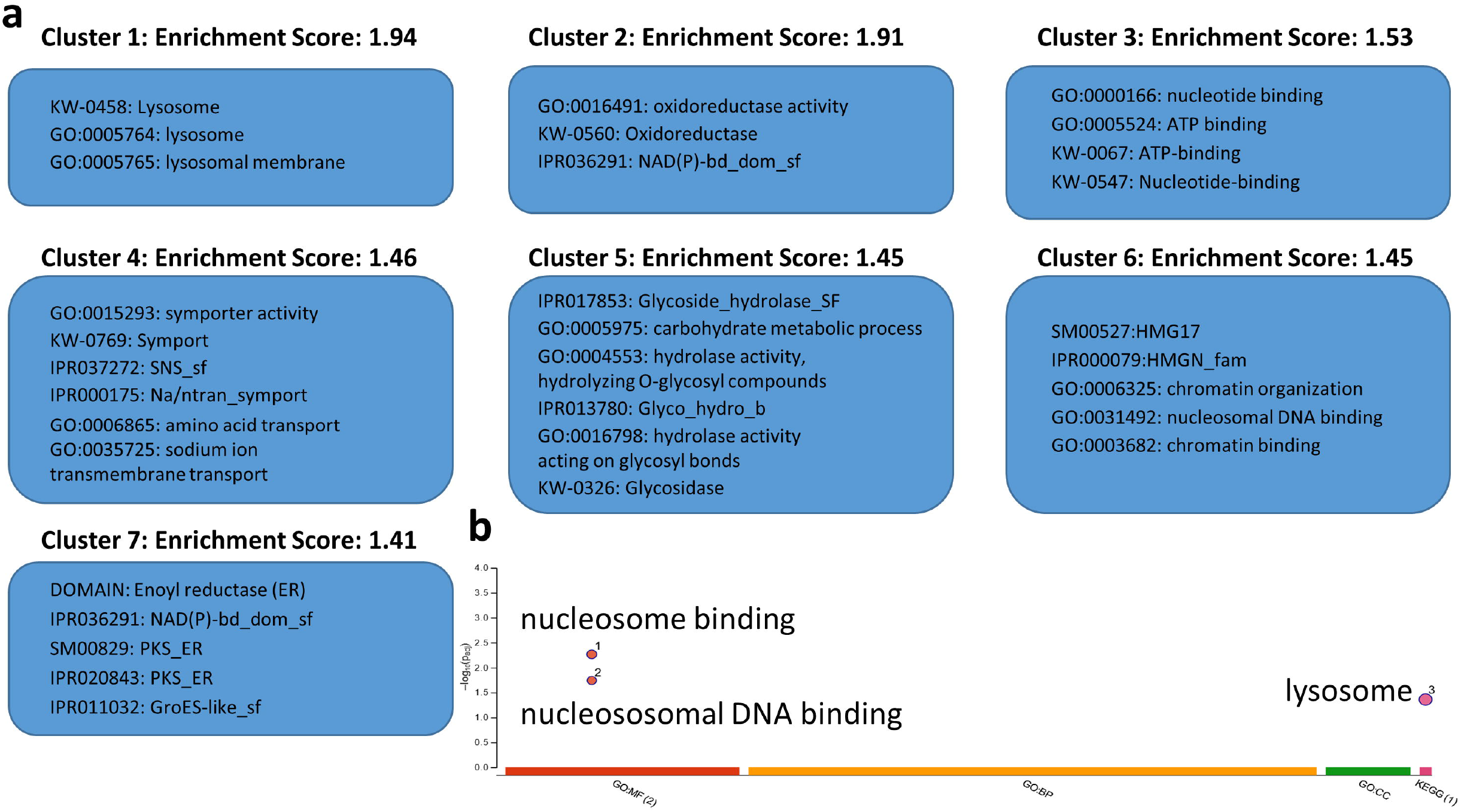
Pathway enrichment analysis of differentially-expressed genes between lrrtm4l1^-/-^ and lrrtm4l1^+/+^ zebrafish. (a) Functional annotation clustering using DAVID pathway analysis revealed seven significantly enriched clusters (enrichment score ≥ 1.3). (b) g:profiler functional enrichment analysis revealed enriched GO and KEGG pathways related to nucleosome binding and lysosome. n=6/group.

## Discussion

Social cognitive impairments are a hallmark of many neurodevelopmental and neuropsychiatric disorders (33). It is unclear why many of these heterogenous disorders present with social abnormalities, but evidence suggests that genetic factors are important regulators of individual sociality. Human genome-wide association studies have identified several loci and genes with a link to aberrant social behaviour, but the function of many of these newly identified genes is currently unknown (19,34). In the current study, we investigated the role of *lrrtm4l1 during* social behaviour, a zebrafish orthologue of the human *LRRTM4* gene that has previously been implicated in autism and children’s aggressive behaviour (17-19,35).

Two major findings within this study suggest that *lrrtm4l1* is an important regulator of zebrafish social behaviour. First, KO zebrafish swim closer together during shoaling assays and second, they spend more time in the vicinity of conspecifics during social interaction tests. This raises the question of what motivates the KO fish to display this altered behaviour. On the one hand, KO zebrafish might want to attack conspecifics to achieve social dominance and therefore the distance between individuals is expected to be smaller. On the other hand, the gene knockout might render them more affiliative towards conspecifics resulting in the same observation. Our data point towards the latter idea, a pro-social phenotype of the mutants, because *lrrtm4l1*^*−/−*^ zebrafish show reduced aggression against their own mirror image and low aggression levels during the social interaction test. This is in line with a previous study which also found reduced aggression in another *lrrtm4l1*^*−/−*^ zebrafish line (Sanger Centre Zebrafish Project, line: sa21708; (26)). It also fits with observations from a study which found that the anti-social and more aggressive *ednraa*^*−/−*^ zebrafish form less cohesive shoals (larger inter-individual distances) and try to attack conspecifics in the social interaction test (36), suggesting an inverse correlation of aggression and shoaling behaviour. An interesting follow-up study would thus be to compare transcriptomic data from *lrrtm4l1*^*−/−*^ and *ednraa*^*−/−*^ zebrafish to identify overlapping or distinct pathways involved in social behaviour regulation.

Another interesting observation of this study is the altered response to diverse social stimuli of mutants compared to WT zebrafish. It is known that environmental context can shape social interactions. For example, many animals are capable of kin recognition, which leads to the preference or avoidance of close relatives depending on the situation (37). Furthermore, social isolation is well known to strongly affect consecutive social interactions (38). In our study, WT zebrafish showed an increase in sociality when exposed to an unknown shoal, an effect absent in mutants. Specifically, *lrrtm4l1*^*−/−*^ zebrafish did not differentiate between known and unknown shoals and spent a large amount of time in the vicinity of either fish shoal. One potential explanation is that mutants do not recognise or remember known conspecifics, an ability that has been demonstrated for WT zebrafish (39,40). Alternatively, it is however also possible that they can distinguish between known and unknown conspecifics, but irrespective of that decide to actively engage in social interactions independent of prior experience. Our data points to the latter, because when we socially-isolated mutant fish, they preferred unknown over known shoals in the same way like WT zebrafish, indicating that under these strongly aversive conditions a distinction between known and unknown is possible. Notable is also the fact that social isolation suppressed the heightened sociality of mutants towards any shoal, indicating that aversive environmental factors can override their genetically-determined pro-social phenotype. This is in line with previous data showing the importance of gene-environment interactions in social dysfunctions (41).

The behavioural phenotype of the *lrrtm4l1*^*−/−*^ mutants was associated with major whole brain transcriptional changes. Of particular interest are the observed expression changes of genes involved in glutamatergic neurotransmission. Previous work in cultured hippocampal neurons and rodents has shown that LRRTM4 is important for the proper functioning of hippocampal excitatory synapses. *Lrrtm4*^*−/−*^ mice show deficits in excitatory neurotransmission in dentate gyrus granular cells and activity-dependent recruiting of AMPA receptors to the postsynaptic membrane (13). In line with this finding, we found that lack of *lrrtm4l1* reduces the expression of *gria2a*, an AMPA receptor subunit, suggesting a conserved effect of LRRTM4 on AMPA receptor signalling across species. The increased expression of *grm2b*, a metabotropic glutamate receptor, suggests that a dysfunction of LRRTM4 signalling has additional effects on glutamatergic neurotransmission, at least in zebrafish. Metabotropic glutamate receptor 2, the protein encoded by the human *GRM2* gene, has been identified as a glutamate auto- receptor regulating glutamate release under high frequency stimulation (42). It has also been studied as a putative target for novel schizophrenia drugs (43), a disorder characterized by major social impairments, which is interesting in the context of the altered social phenotype of the *lrrtm4l1* mutants.

Another interesting finding of this study is the observed changes in genes of the kynurenine pathway, indicating that lack of *lrrtm4l1* leads to changes in kynurenine metabolism. Kynurenine is one of the metabolites of tryptophan, besides the neurotransmitter serotonin and the microbially-derived indoles. It is further metabolized into important neuroprotective, but also neurotoxic, metabolites such as kynurenic acid and quinolonic acid (QUIN), respectively (44). As described above, the altered expression of *tdo2a* and *kmo*, two important enzymes in the kynurenine pathway, is expected to have at least two major effects. First, the lack of tdo2a, the rate limiting enzyme of the kynurenine pathway, in mutant zebrafish leads to a higher availability of tryptophan for other pathways, such as the serotonergic pathway. In the brain this may lead to manifold effects depending on the brain area and receptor activated. Of interest in relation to social behaviour are findings that serotonin positively correlates with social behaviour and sensitivity to social factors (45). Second, the higher expression of *kmo* which converts kynurenine to 3-hydroxy kynurenine (3-HK), may lead to an accumulation of 3-HK, another known neurotoxic compound of the kynurenine pathway, and the neurotoxic QUIN further downstream in the pathway (44). Elevated 3-HK and QUIN levels have been implicated in the pathology of multiple neurodegenerative diseases (46), but their role in social abnormalities is currently unclear.

The transcriptomic data also revealed expression changes of genes implicated in synaptic plasticity. Specifically, we detected a downregulation of *sypa* in *lrrtm4l1* mutants, which is the zebrafish orthologue of human *synaptophysin*. Synaptophysin is a major protein of synaptic vesicle membranes and therefore an important component of synaptic neurotransmission in all major neurotransmitter systems (47). Although evidence about the role of synaptophysin during social behaviour is sparse, a recent study found that social defeat stress leads to reduced synaptophysin levels in the medial prefrontal cortex of Syrian hamsters (48). This suggests a role of synaptophysin in adaptation or coping behaviour after social stressors. Given the presence of synaptophysin in synaptic vesicles, reduced synaptophysin might be an indicator of a reduction in presynaptic terminals. Like *sypa, glypican 2 (gpc2)* was also downregulated in the mutants. This is of interest, because glypican2 belongs to the glypican protein family that has been identified as presynaptic binding partners of LRRTM4 (9). Therefore, the downregulated expression of glypican2 in *lrrtm4l1* mutants might suggest a compensatory response and a similar transsynaptic signalling of *lrrtm4l1* in zebrafish compared to its mammalian orthologue. In contrast to *sypa* and *gpc2, sema3f* was upregulated in mutant zebrafish. This gene encodes an axonal guidance protein that binds to its receptor of the plexin protein family. Semaphorin-plexin signalling in general is known to be important for numerous brain plasticity processes, such as neurogenesis and neuronal cell migration (49). Interestingly, it has been shown that conditional knockout of semaphorin F in GABAergic interneurons of mice leads to social deficits (50). This in line with the upregulated *sema3F* expression in *lrrtm4l1*^*−/−*^ zebrafish that show a pro-social phenotype. Finally, an unexpected finding of this study is the enrichment of lysosomal-associated genes in the transcriptomic dataset. To our knowledge this is the first description of a potential link between LRRTMs and lysosomal processes. There is currently no evidence of *LRRTM4* expression in lysosomal membranes or of signalling pathways connecting LRRTMs in the postsynaptic membrane with the lysosome. Therefore, this finding warrants further investigation about the relevance of this potential interaction for neuronal function and behaviour.

A limitation of this study is that we do not know the exact neuronal circuits and cell types underlying the phenotype and gene expression changes observed in this study. However, it has recently been shown that *lrrtm4l1* has a brain region-specific expression pattern (26), suggesting that the lack of lrrtm4l1 protein in brain areas with high *lrrtm4l1* expression is important for the observed behavioural changes in this study. The brain areas with the highest *lrrtm4l1* expression in adult zebrafish brain are the anterior telencephalon and the inferior lobe (26), that have been implicated in numerous higher brain functions and multisensory processing, respectively (51,52). The high *lrrtm4l1* expression in the telencephalon is of particularly interest for the current study, given that many nuclei of this region belong to the social decision-making network (SDMN). The SDMN is well known to govern social behaviour across species and activity changes in the network have major effects on pro- or anti-social behaviour (53,54). In zebrafish, social interactions alter the functional connectivity in the SDMN (55), a process that might be impaired in *lrrtm4l1* mutants. Future studies employing single-cell RNAseq of the telencephalon or advanced genome engineering techniques to generate neuron-specific conditional *lrrtm4l1* knockout animals would be helpful to gain deeper mechanistic insights. Finally, it is unclear, whether other zebrafish orthologues of *LRRTM4* such as *lrrtm4l2* play a role during social behaviour as well and whether these orthologues compensated some of the effects induced by *lrrtm4l1* deletion.

### Conclusion

In conclusion this study suggests that *lrrtm4l1* is an important regulator of social behaviour in zebrafish. Germ-line knockout of *lrrtm4l1* leads to a pro-social phenotype of mutant zebrafish, which is accompanied by alterations of key genes in neurotransmitter signalling and synaptic plasticity. In a translational view, these findings suggest LRRTM4 as a potential therapeutic target for conditions like anti-social personality disorder or as a potential pathophysiological factor contributing to disorders characterized by hypersociability such as Williams syndrome. Further studies are needed to elucidate whether manipulation of LRRTM4 signalling is also socially beneficial in other model organisms and in the context of disease models characterized by social impairments.

## Methods

### Experimental animals and housing conditions

Experiments were performed with mixed-sex, age-matched (10-months) adult *lrrtm4l1*^*−/−*^ and corresponding AB wild-type (*lrrtm4l1*^+/+^) zebrafish. Animals used for experiments were obtained from multiple crosses and tanks to avoid batch effects. Mixed zebrafish groups (50:50 - male:female ratio) were used to avoid male animal bias and behavioural experiments were performed in the morning between 9 and 12am to avoid diurnal variations. All animals were kept at the same stocking density of 5-6 adult/fish per litre to ensure consistent environmental conditions. The zebrafish facility at the University of Portsmouth features optimal housing conditions for zebrafish, housed in a recirculating water system (Aquaneering, Fairfield Conrolec Ltd., Grimsby, UK), a 14-hour light/10-hour dark cycle with lights turned on at 08:30 am, standard feeding schedules (from 5 days post fertilization [dpf], larvae were fed ZM-100 fry food (Zebrafish Management Ltd., Winchester, UK) until adulthood, where they were transitioned to flake food and live brine shrimp 4/day [1/day on weekends]), regulated water temperature (25-27°C), pH of 8.4 (± 0.4) and dedicated personnel. Animal experiments were approved by a local Animal Welfare and Ethical Review Body (AWERB) and conducted under the UK Home Office project (PP8708123) and personal licences.

### Generation of lrrtm4l1 knock out zebrafish using CRISPR/Cas9

*lrrtm4l1* mutant zebrafish were generated using CRISPR/Cas9 following a previously described protocol (56). Briefly, AB wild-type zebrafish embryos were microinjected with a mix of single guide RNA (sgRNA) targeting an early part of exon 2 of the gene (target sequence: 5’-GGAAGGCAGACGACTCACAGTGG-3’) as predicted by CHOPCHOP software (57) and Cas9 mRNA. The sgRNA was generated by in vitro transcription using the mMESSAGE mMACHINE kit (Life Technologies) on a sgRNA expression vector (pDR274; Addgene plasmid #42250) as a template. Cas9 mRNA was transcribed from the pMLM3613 vector (Addgene plasmid #42251) using the mMESSAGE mMACHINE T7 Ultra Kit (Life Technologies). After cleanup, RNA concentrations were quantified, mRNAs were diluted and stored at -80°C until injection. One-cell embryos were co-injected with approximately 1 nl total volume of Cas9-encoding mRNA (216 ng/μl) and sgRNA (36 ng/μl). Injection efficiency was assessed by amplifying the target region by fluorescence PCR and screening for mutations using capillary gel electrophoresis following the protocol of Varshney et al. 2017 (58). Germ-line transmitting founders were identified by outcrossing injected individuals (F0 generation) to wild-type zebrafish and screening the offspring (F1 generation) for mutations at the target site. F1 zebrafish heterozygous for a 1 bp insertion in exon 2 of the gene (*lrrtm4l1*^*uol3*^) were then in-crossed and genotyped by capillary gel electrophoresis to obtain homozygous mutants (*lrrtm4l1*^*−/−*^) and corresponding wild-type fish (*lrrtm4l1*^*+/+*^) used for breeding of animals for further experiments.

### Shoaling behaviour

To analyse shoaling behaviour, five adult fish per genotype were placed together in a white tank (150 w x 200 d x 150 h [mm]) within a Zantiks (Zantiks Ltd., Cambridge, UK) AD unit and videorecorded for 30 min as previously described (59) Animal tracking (X and Y coordinates) was achieved using the multi-animal mode of DeepLabCut (60), with key points defined at the head, trunk, and tail. Using ResNet 101, we trained a deep neural network on 90 frames across both groups, followed by iterative refinement through outlier correction and the addition of 30 frames. The centre point of each fish was used to determine minimum, maximum, and average inter-fish distances, as well as the shoal area. These parameters were quantified using an R script designed to precisely analyse shoaling behaviour throughout the test.

### Social interaction test

To analyse social behaviour, we used a modified version of the visually-mediated social preference test (VMSP; (22)). In this assay, a test fish was placed in the central compartment of a 5-chamber tank (dimensions: 360 w x 270 d x 300 h [mm]) for 30 min. The smaller surrounding compartments cannot be accessed by the test fish, but small holes in the separating transparent walls permit exchange of water and cues. To analyse social behaviour, one of the smaller compartments was filled with a group of four stimulus fish and the time that the test fish spends in the near of the stimulus fish (social interaction zone within the central compartment; Fig.1c) was quantified. To analyse the effect of social context on the behaviour of the test fish, the test fish was exposed to known or unknown stimulus shoals or isolated for 2 hours before testing or both. The behaviour of the test fish was recorded and video-tracked using a Zantiks LT unit and Zantiks videotracking software.

### Aggressive behaviour

The mirror-induced aggression (MIA) paradigm was used to record the fish’s interaction with its mirror image and to score the amount of time spent in agonistic behaviour (61). For this, individual fish were placed in a glass tank (15 × 10 × 30 cm), where the sides and back wall were covered with grey foam rubber. A mirror was placed in front of the tank at an angle of 22.5° and the test fish was allowed to explore the arena freely for 5 min. Videos were recorded from the top using FlyCap2 software (Point Grey Research; Canada) and the time spent in aggressive display against the mirror image was evaluated manually by an experienced investigator blinded for the experimental groups, as previously described (61). During the VMSP aggressive behaviour against the transparent wall separating the test fish from the stimulus fish was also quantified in a similar manner. Specifically, three time bins of the 30-min testing period (seconds: 240-360; 840-960; 1440-1560) were arbitrarily selected for manual aggression analysis (in total 6 min/stimulus fish) in order to quantify the average time each stimulus fish spent in aggressive display against the known or unknown shoal.

### RNAseq and bioinformatics

For the RNAseq experiment, zebrafish were euthanised in ice-cold water (4°C) and brains were extracted in ice-cold PBS. Brains were frozen on dry ice and stored at -80°C until further processing. RNA was extracted using Trizol and phenol/chloroform extraction according to standard operating procedures at BGI Genomics (Shenzen, China). RNA samples were then used for mRNA library preparation and sequenced by paired-end (2×100bp) sequencing on an DNBseq-G400 sequencing platform at BGI Genomics. The generated paired-end raw sequence data were analysed with a Galaxy pipeline as previously described (61). Briefly, raw sequence data were quality controlled (Fast QC; Galaxy Version 0.11.9) and sequencing adapters as well as reads shorter than 50 base pairs were removed with Trim Galore! (Galaxy Version 0.6.3.) to increase the mapping quality. On average 81.8% (stdev 0.004%) of the reads could be successfully uniquely mapped with the RNAStar aligner (((62); Galaxy Version 2.7.8a) to the zebrafish reference genome GRCz11. Final transcript count data was generated with the HTSeq framework ((63); Galaxy Version 0.9.1) for high throughput sequencing data using default settings. All analysis were conducted on a private Galaxy instance running on the MedBioNode cluster from the Medical University Graz. Further downstream analysis was conducted with the statistical program R version 4.4.1 within the free RStudio environment. Differential gene expression analysis was performed with DESeq2 (version 1.44.0; (64)) on the count table as output from HTSeq framework. The Database for Annotation, Visualization and Integrated Discovery (DAVID; v2024q2; (31)) was used for pathway enrichment analysis by clustering DEGs and associated biological annotation terms into functional groups. Enrichment score cutoff in DAVID was set to 1.3, which corresponds to a corrected p value of 0.05. In addition, g:profiler (32) was used for functional enrichment analysis of DEGs.

### Statistics

Statistical analysis was performed with the Prism 10.2.2 (Systat Software Inc, San Jose, USA) software package. Student’s t test (two-tailed) was used to compare data of two normally distributed groups, while Mann-Whitney-U test was used to compare data of two not-normally distributed groups. For experiments involving analysis over time, we conducted a Linear Mixed Effects model using SPSS with group (WT vs. KO /known/unknown and/or /isolation), time bin (1–30 min), and their interaction as fixed effects. Subject ID was included as a random effect with a Variance Components (VC) covariance structure, which assumes independent variance for each subject within the social interaction tests. Graphpad Prism software was then used to run unpaired Student’s t-tests for significance assessments. P-values < 0.05 were considered as statistically significant.

## Acknowledgments

We are grateful to the animal care takers at the University of Leicester and the University of Portsmouth for zebrafish care and maintenance. This research was funded fully or in part by the Austrian Science Fund (FWF) [10.55776/PAT5589324] and The Fisheries Society of the British Isles (FSBI, Postdoctoral International Travelling Fellowships PITF). For open access purposes, the author has applied a CC BY public copyright license to any author accepted manuscript version arising from this submission. CH was funded by Dstl (MOD, UK). GP is funded by the FWF within the PhD program Molecular Medicine of the Medical University of Graz.

