## Supplementary information for "Genetic disruption of leucine rich repeat transmembrane protein 4 like 1 induces a pro-social behavioural phenotype in zebrafish"

*Corresponding authors:

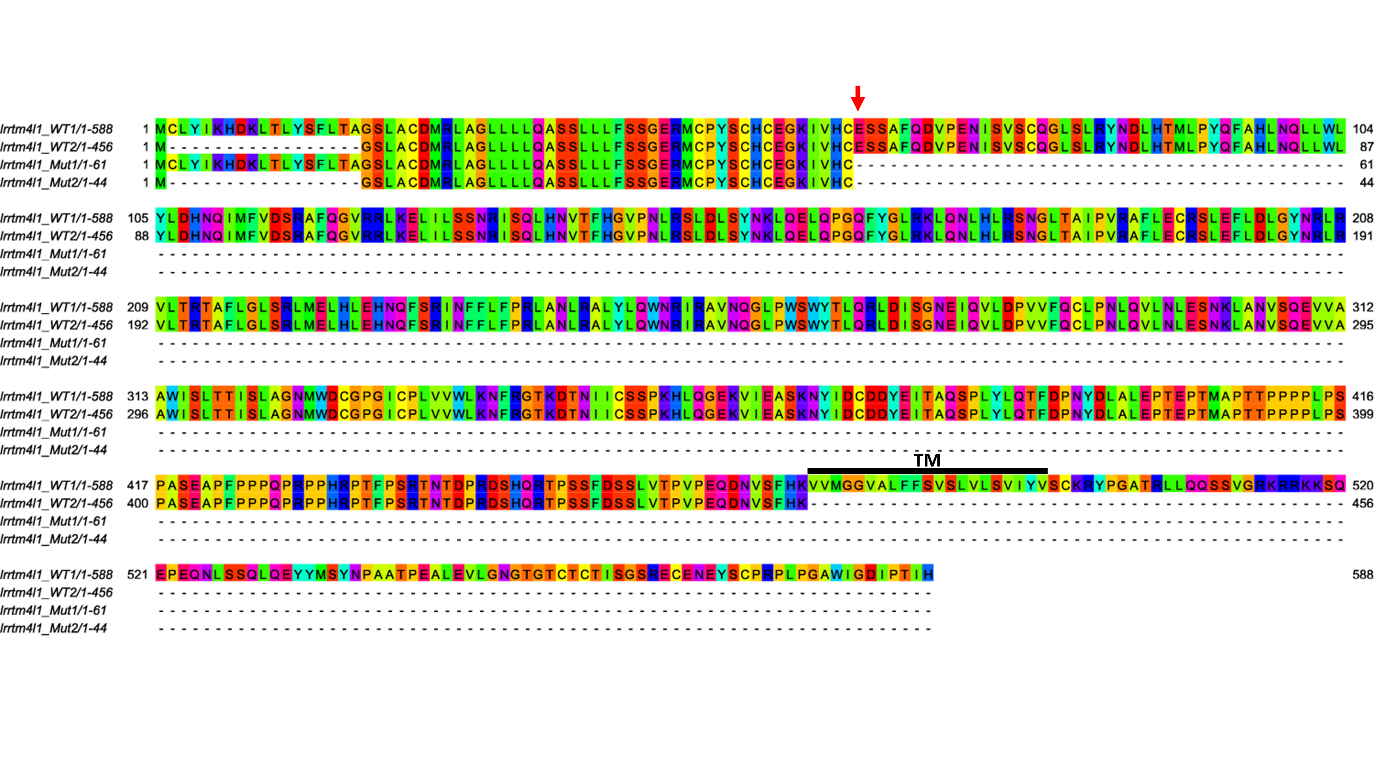

***Supplementary Figure 1. Predicted amino acid sequences of the leucine rich repeat transmembrane neuronal 4 like 1 (lrrtm4l1) mutant proteins.*** *Wild-type (lrrtm4l1_WT1, Ensembl transcript ID: ENSDART00000113737.4 and lrrtm4l1_WT2, Ensembl transcript ID: ENSDART00000141364.2) and mutant (lrrtm4l1_Mut1 and lrrtm4l1_Mut2) proteins have a conserved amino acid sequence until amino acid position 61 (indicated by a red arrow), where a 1 bp insertion in the mutant allele leads to a loss of reading frame and an early stop codon. The transmembrane domain was predicted using TMHMM Server v. 2.0 and is highlighted by a black line. The amino acid residues were coloured with the Taylor colour scheme using the open-source Jalview software (1).*

**References**

(1) Waterhouse AM, Procter JB, Martin DM, Clamp M, Barton GJ. Jalview Version 2--a

multiple sequence alignment editor and analysis workbench. Bioinformatics 2009 May

1;25(9):1189-1191.
